# Evolutionary Based Adaptive Dosing Algorithms: Beware the Cost of Cumulative Risk

**DOI:** 10.1101/2020.06.23.167056

**Authors:** Hitesh B. Mistry

## Abstract

Application of theories from ecology to cancer is growing. One such idea involves using drug sensitive cells to control drug resistant cells by cycling treatment on and off based on a marker of tumour burden. Many literature studies have highlighted the benefit of this approach when using time till reaching a certain level of burden as an end-point. These studies though have not considered that patients need to survive up until a certain time-point with a higher level of tumour burden compared to standard dosing to gain this benefit. Within this study once this cumulative cost is accounted for it can be seen that adaptive dosing, counter-intuitively, is likely to lead to poorer prognosis than continuous dosing. This study highlights that evolutionary based adaptive dosing algorithms may not be the “parachute” its protagonists believe it to be.

---

There is growing interest in applying concepts from ecology to cancer treatments as we begin to see cancer cells as part of a complex ecosystem.^1^ One concept from ecology which is gaining traction is the use of predator-prey (Lotka-Volterra^2,3^) models to design novel dosing strategies to control the tumour.^4,5^ Before considering its application in cancer, it is important to introduce it from an ecologist’s perspective.

Consider two species foxes (predator) and rabbits (prey). The number of foxes will depend on the number of rabbits and vice versa. As foxes eat rabbits and reproduce the rabbit population will decrease if rabbits reproduce at the same rate as they have always done. At some point the number of foxes will decrease as there isn’t enough food to sustain the population. As foxes decrease the rabbit population starts to increase again. This leads to cyclical levels of rabbits and foxes with a certain time delay. The temporal oscillations of rabbits or foxes though have little effect on say the forest they reside in or the planet the forest is on. The key point here is that there is no risk to the forest or the planet when we first observe the system or over time from that first observation. On the other hand, when a cancer patient is about to be treated the reason for choosing to treat is that there is a heightened risk of an adverse outcome (e.g. death or cancer progression) if the disease is left untreated. The above predator/prey example in ecology therefore has a different connotation in cancer when cumulative risk is considered as we now discuss.

The simplest predator-prey setup in cancer is as follows. The predator and prey are drug sensitive and drug resistant cancer cells, respectively. The drug sensitive cells are assumed to be in excess of the resistant cells and outcompete the resistant cells for resource. The system at the time of observation is such that if left unperturbed will simply lead to an expansion of both the drug sensitive and resistant cells. The adaptive dosing algorithm based on evolutionary concepts simply involves setting disease burden threshold levels to decide when to stop and start treatment.

Unlike the ecology example the total number of predator-prey species, cancer cells, are causing distress to the patient. Therefore, in cancer both the current and historical level of tumour burden directly relates to the probability of that patient overcoming the resultant adversity from cancer (e.g. survival). This has been observed in many biostatistical analyses which have found tumour burden to be a time-dependent covariate for patient survival.^6–10^ The time-series generated from a Lotka-Volterra model has never been explicitly analysed within the context of these statistical survival models. In this short report we consider the consequence of adaptive dosing therapy algorithms in the context of cumulative risk.

Let us consider the simplest version of a Lotka-Volterra model in the context of cancer,

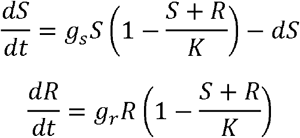

The rate of change of the drug sensitive cells, *S*, is equal to the difference between the growth rate *g_s_* and the loss due to drug effect, *d*, and the competition from the resistant cells, *R* at time t. The rate of change of the drug resistant cells is equal to the difference between the growth rate, *g_r_* and the competition from drug sensitive cells at time *t*. For simplicity we assumed the competition is similar between drug sensitive and resistant cells and the carrying capacity, *K*, was also set to 1. If competition is not considered the above model reduces to a simple exponential decay/growth model, which have been applied to many clinical trial data-sets.

The adaptive therapy strategy has predominantly centered around targeted therapy and so we shall use a set of parameter values, based on fitting an exponential decay/growth model to 100s of patients, for the targeted agent, Gefitinib, see Table 1 (Table S7 in Mistry et al. 2019^11^). A net decay rate *g_s_* – *d* was estimated in Mistry et al. 2019.^11^ In order to estimate, *g_s_* and thus *d*, we shall use the following argument. The level of tumour burden is typically assessed 6-8 weeks after treatment initiation in most clinical trials of targeted agents.^11^ If we take the latter time-point, 8 weeks, as the first time that a tumour is assessed after treatment initiation and assume the patients’ disease progressed according to RECIST^12^ criteria on that visit i.e. a minimum 20% increase in tumour size is observed compared to pre-treatment measurement. Then a lower bound for *g_s_* can be derived and thus a value for *d*, see Table 1. The proportion of tumour that is drug sensitive versus resistant is also taken from Mistry et al. 2019. Initial value of the total tumour burden was chosen to be 0.75, 75% of the total carrying capacity. This value was chosen to provide a reasonable depth of tumour shrinkage under continuous treatment to explore adaptive dosing regimens.

**Table 1:** Parameter values

| Parameter Description | Parameter | Value | Reference |
| --- | --- | --- | --- |
| Proportion of tumour drug sensitive | $S(0)/R(0)$ | 0.73 | Table S7 Mistry et al. 2019<sup>11</sup> |
| Initial Tumour Burden | $S(0) + R(0)$ | 0.5 | This study |
| Net sensitive clone loss rate | $d' = g_s - d$ | 0.0166 per day | Table S7 Mistry et al. 2019<sup>11</sup> |
| Resistant clone growth rate | $g_r$ | 0.0017 per day | Table S7 Mistry et al. 2019<sup>11</sup> |
| Sensitive clone growth rate | $g_s$ | $\log(1.2)/56 = 0.0033$ per day | This study |
| Sensitive clone drug induced decay rate | $d$ | 0.0133 per day | This study |
| Event rate | $a_0$ | 0.003 per day | This study |

The adaptive therapy algorithm that has been discussed in the literature involves the patient continuing treatment until the tumour has decreased by X% compared to pre-treatment levels.^13^ At that point the treatment is stopped and re-started again once the disease has reached pretreatment levels. This cycling of treatment continues until the patient no longer achieves a X% fall from pre-treatment values. This can be mimicked in the model by turning the decay rate *d*, on/off according to the level of tumour burden. Let us now consider the dynamics of the system under notreatment, continuous and adaptive treatment.

Under the control situation i.e. no treatment the total tumour burden is given by the red-line in Figure 1. Upon the invention of a new treatment which is given as a daily continuous therapy the dynamics of the system follows the blue line in Figure 1. We see the tumour regress before resistance emerges. Based on that simulation the threshold value chosen for adaptive therapy treatment discontinuation was chosen to be 30% reduction in tumour burden compared to baseline. The therapy is then re-started once the tumour reaches the baseline value again. Using a 30% threshold for treatment discontinuation and 0% for treatment initiation leads to the green line simulation in Figure 1. We can see that adaptive therapy gives control i.e. the time till adaptive therapy reaches maximal burden is longer than that for continuous therapy. Therefore, adaptive dosing appears to be a success. However, we have not taken into consideration that the benefit is conditional on surviving up until the time where the green line intersects with the blue line i.e. continuous therapy equals adaptive therapy. We shall now consider how this conditional statement effects our conclusion about the effectiveness of adaptive therapy.

**Figure 1:**
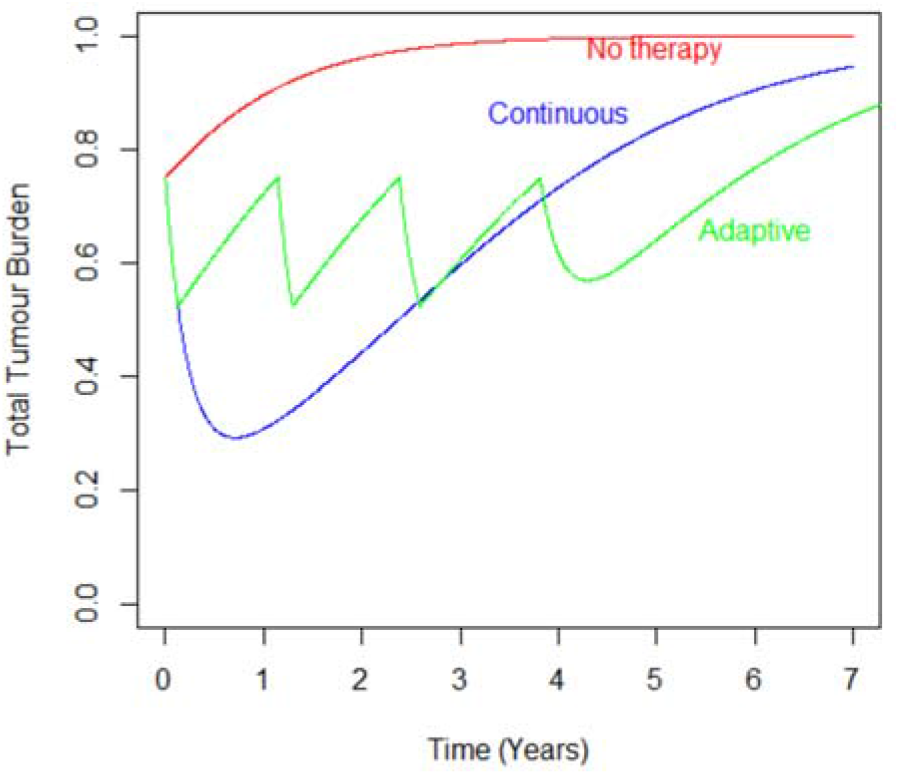
Plot showing the temporal evolution of the total species (S+R) under no therapy (red line), continuous (blue line) or adaptive (green line) drug-treatment.

In order to explore whether a patient is likely to survive long enough to benefit from adaptive therapy we shall link the dynamics of total tumour burden to a simple survival model, based on the exponential distribution. The expected survival probability at a given point in time is given by,

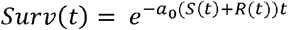

A value for *a*_0_, see Table 1, was chosen such that without therapy the median survival probability was 12 months. The survival probability over time of no-treatment, continuous and adaptive therapy, using the thresholds described above, can be seen in Figure 2. The plot shows that the new treatment, given continuously, increases survival over no therapy but that choosing to use an adaptive dosing algorithm gives less benefit compared to continuous dosing.

**Figure 2:**
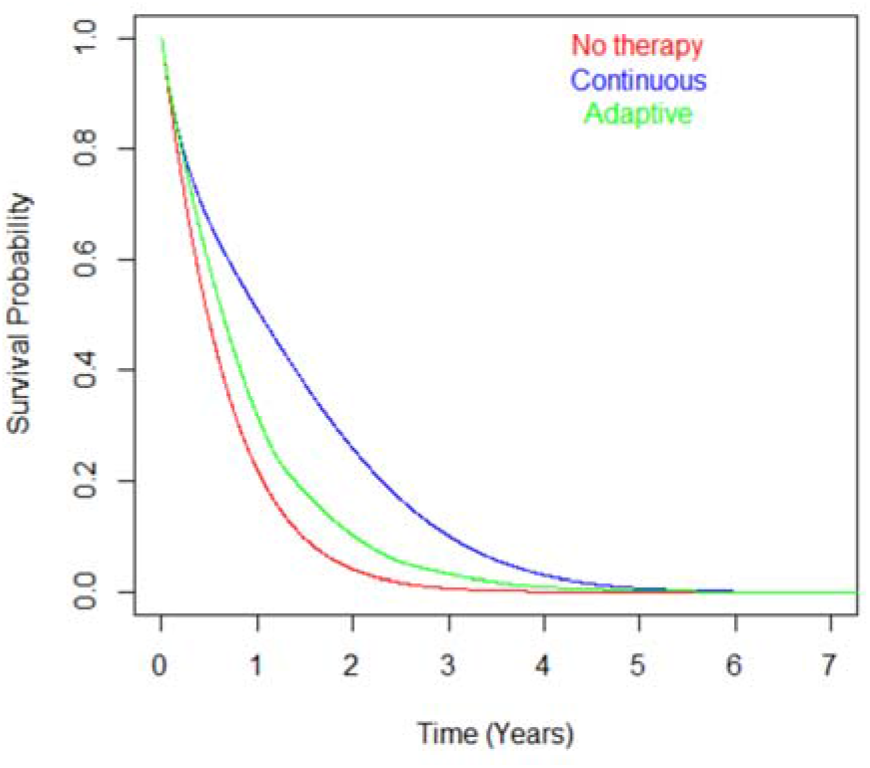
Plot showing the temporal evolution of the survival probability under no therapy, continuous drug-treatment or adaptive.

The reason for the above result is that continuous therapy reduces risk dramatically by eradicating as much of the tumour as possible as quickly as possible, whereas adaptive therapy does not, see Figure 1. Adaptive therapy only modestly reduces a patient’s risk of experiencing an event and keeps the patient in that state until the resistant clone emerges. The more we try to control the tumour the closer our strategy equates to giving no treatment at all, see Figure 3. In fact, the best scenario is to give as much treatment as possible as quickly as possible, the current paradigm in drug development. These results argue that adaptive therapy may not be the “parachute” the community devising these strategies seem to believe.^14^ No preclinical system, in-vitro or in-vivo, contains the concept of cumulative risk like the human system. Therefore, human studies and in particular randomized control trials are the only option to truly assess whether adaptive therapy is likely to lead to harm or benefit.

**Figure 3:**
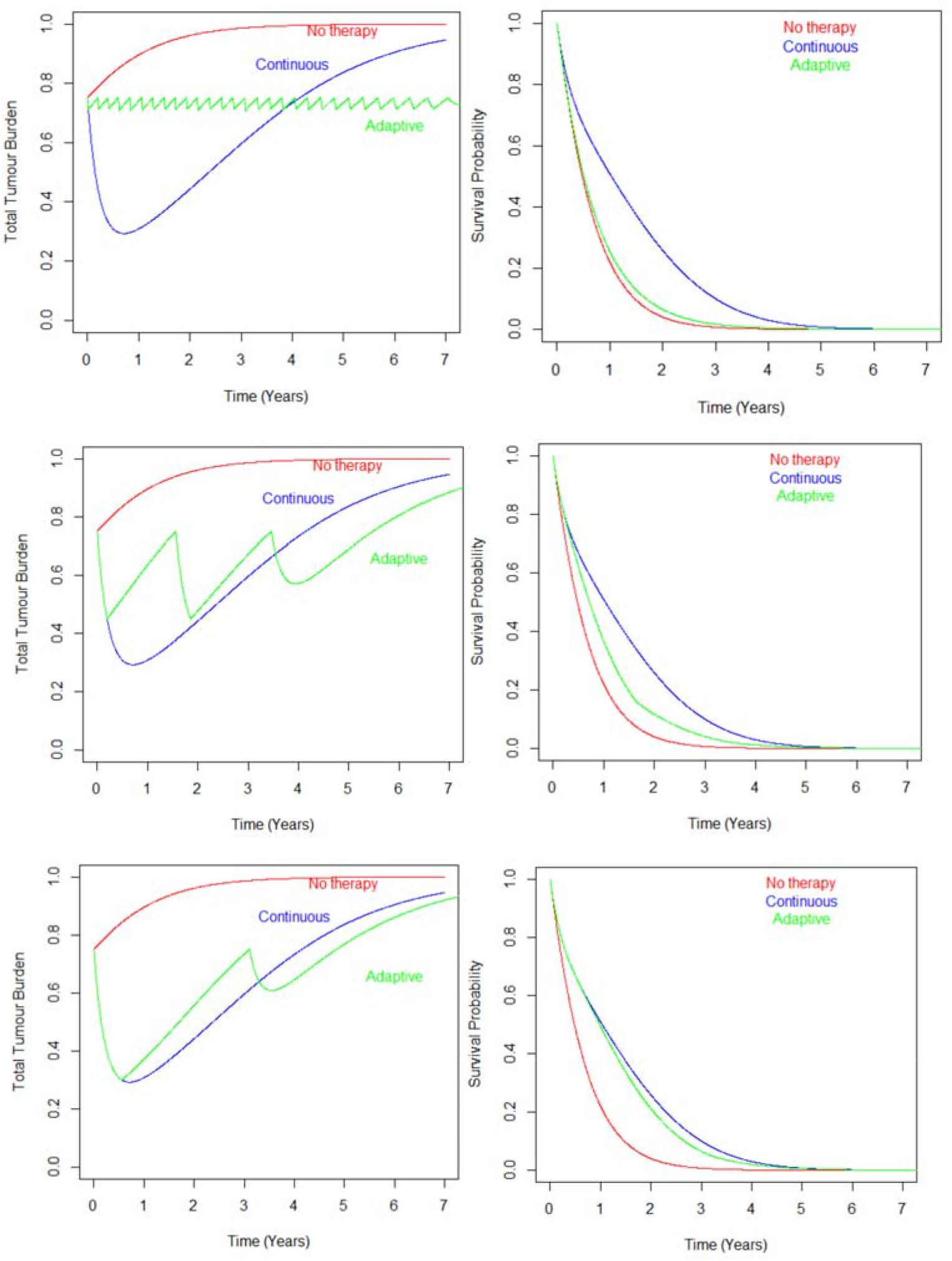
Plot showing the temporal evolution of the total tumour burden and corresponding evolution of survival probability under no therapy, continuous drug-treatment or adaptive with different adaptive therapy treatment discontinuation thresholds, 5% (top), 40% (middle) and 60% (bottom) reduction.

